# Matrix-targeted Nanoparticles for MMP13 RNA Interference Blocks Post-Traumatic Osteoarthritis

**DOI:** 10.1101/2020.01.30.925321

**Authors:** Sean K Bedingfield, Fang Yu, Danielle D. Liu, Meredith A. Jackson, Lauren E. Himmel, Hongsik Cho, Juan M. Colazo, Leslie J. Crofford, Karen A. Hasty, Craig L. Duvall

## Abstract

Osteoarthritis (OA) is a debilitating and prevalent chronic disease, but there are no approved disease modifying OA drugs (DMOADs), only pharmaceuticals for pain management. OA progression, particularly for post-traumatic osteoarthritis (PTOA), is associated with inflammation and enzymatic degradation of the extracellular matrix. In particular, Matrix Metalloproteinase 13 (MMP13) breaks down collagen type 2 (CII), a key structural component of cartilage extracellular matrix, and consequently, matrix degradation fragments perpetuate inflammation and a degenerative cycle that leads to progressive joint pathology. Here, we tested targeted delivery of endosome-escaping, MMP13 RNA interference (RNAi) nanoparticles (NPs) as a DMOAD. The new targeting approach pursued here deviates from the convention of targeting specific cell types (*e.g*., through cell surface receptors) and instead leverages a monoclonal antibody (mAbCII) that targets extracellular CII that becomes uniquely accessible at early OA focal defects. Targeted mAbCII-siNPs create an *in situ* NP depot for retention and potent activity within OA joints. The mAbCII-siNPs loaded with MMP13 siRNA (mAbCII-siNP/siMMP13) potently suppressed MMP13 expression (95% silencing) in TNFα-stimulated chondrocytes *in vitro*, and the targeted mAbCII-siNPs had higher binding to trypsin-damaged porcine cartilage than untargeted control NPs. In an acute mechanical injury mouse model of PTOA, mAbCII-siNP/siMMP13 achieved 80% reduction in MMP13 expression (p = 0.00231), whereas a non-targeted control achieved only 55% silencing. In a more severe, PTOA model, weekly mAbCII-siNP/siMMP13 long-term treatment provided significant protection of cartilage integrity (0.45+/− .3 vs 1.6+/−.5 on the OARSI scale; p=0.0166), and overall joint structure (1.3+/−.6 vs 2.8+/−.2 on the Degenerative Joint Disease scale; p<0.05). Intra-articular mAbCII-siNPs better protected articular cartilage (OARSI score) relative to either single or weekly treatment with the clinical gold stand steroid treatment methylprednisolone. Finally, multiplexed gene expression analysis of 254 inflammation-related genes showed that MMP13 inhibition suppressed clusters of genes associated with tissue restructuring, angiogenesis (associated with synovial inflammation and thickening), innate immune response, and proteolysis. This work establishes the new concept of targeting unique local extracellular matrix signatures to sustain retention and increase delivery efficacy of biologics with intracellular activity and also validates the promise of MMP13 RNAi as a DMOAD in a clinically-relevant therapeutic context.

**Abstract Figure:**
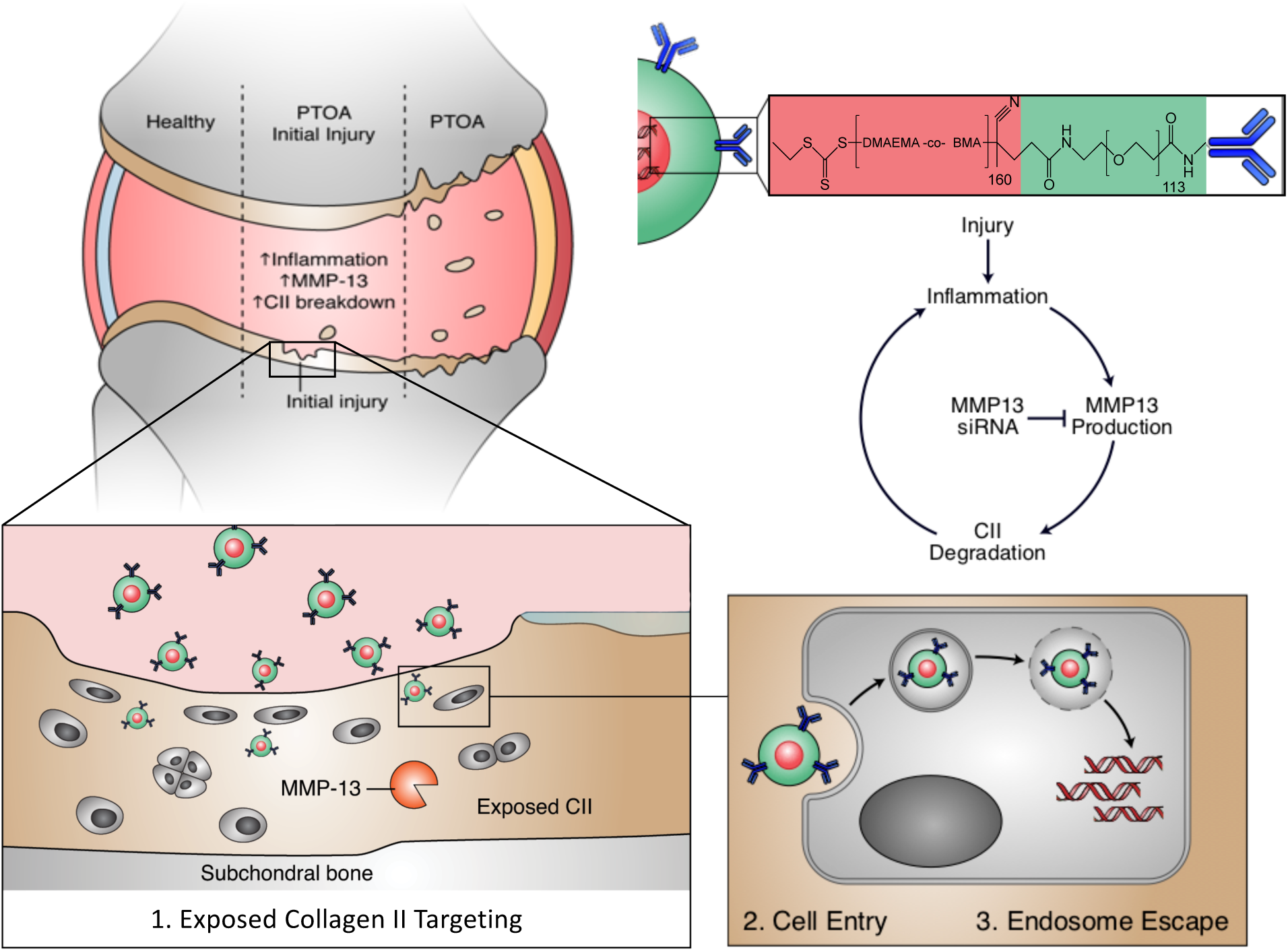
PTOA targeted delivery of MMP13 siRNA to block disease progression. The top left schematic illustrates the progression (left to right) from healthy knee joint, through inflammation induction following traumatic injury, to cartilage loss and degenerative joint disease (including synovial response). Degradation of cartilage enhances inflammation, inducing a degenerative cycle (middle right). The bottom of the graphic illustrates the concept of the matrix targeted nanocarriers for enhanced retention and activity of MMP13 siRNA at sites of cartilage injury.

## Introduction

Osteoarthritis (OA) is a chronic degenerative disease affecting joints and their surrounding tissues. Initial symptoms include pain and loss of mobility, resulting in significant loss in quality of life. The burden of OA on the United States healthcare system is estimated to be over $44 billion annually in direct costs and up to double that in indirect economic burden (1). More than 25% of the American population over the age of 45 is afflicted with OA, and its prevalence is anticipated to increase over the next twenty years (2).

OA is a disease of the entire joint that involves a complex interplay between mechanical and biochemical factors (3). Some well-established risk factors include poor joint alignment or injury (4), obesity (5), genetic disposition (6), and aging (7). Multiple signaling molecules are known to be central to OA pathogenesis such as interleukin (IL-1β), Wnt, c-Jun N-terminal kinase (JNK), and reactive oxygen species (ROS) (3, 8, 9). All of these signaling pathways independently converge toward increased production of matrix metalloproteinases (MMPs), a step of critical importance in cartilage degradation and progression of OA symptoms (3).

Post-traumatic osteoarthritis (PTOA) is a form of OA induced by a mechanical joint injury. Common injuries include ligament and meniscal tears, cartilage damage, bone fractures from high impact landings, and dislocations. These injuries are particularly common among young athletes and military personnel and result in an accelerated pathology, requiring surgical intervention 7-9 years earlier on average than standard OA (10). Approximately 12% of all OA cases are PTOA in the United States, and PTOA tends to have greater cost relative to normal OA both in annual expenditure (∼$3 billion) and in quality adjusted life years (QALYs) due to its younger age of onset and accelerated progression (11). Following a PTOA-initiating injury, the mechanical disturbance of the extracellular matrix (ECM) stimulates synoviocytes and chondrocytes to produce inflammatory cytokines and MMPs (12). MMP13 is an enzyme able to catalytically degrade collagen II (CII), a key cartilage structural component. Degradation of CII disrupts chondrocytes by destroying the ECM in their surroundings and also releases soluble ECM degradation biproducts which have inflammatory signaling properties that trigger the OA degenerative cycle (3, 13). Inflammatory activation of the full joint and surrounding tissues (synovium and subchondral bone) perpetuates this cycle until cartilage destruction is complete. Because PTOA commonly has a defined and predictable initiating event, there is potential for early therapeutic intervention to block disease onset or progression at an early stage.

Current pharmaceutical management of OA is solely palliative, and no disease modifying OA drugs (DMOADs) are clinically approved. There are five FDA-approved corticosteroids for intra-articular OA therapy, but these therapies provide only temporary pain relief. Steroids do not target the underlying cause of disease and are not recommended for long-term management(14), as they have been shown to actually cause cartilage volume loss (when given 4 times per year for 2 years)(15) and have associations with chondrotoxicity (16). MMP13 is a key proteolytic driver of cartilage loss in OA as indicated by reduced OA progression in surgically induced OA in MMP13 knockout mice and in wild type mice treated with broad MMP inhibitors (17). Unfortunately, clinical trials on MMP small molecules inhibitors (tested mostly for cancer treatment) have been suspended due to pain associated with musculoskeletal syndrome (MSS). Patient MSS is believed to be linked to systemic delivery of small molecules that non-selectively inhibit multiple MMPs, some of which (MMP2, 3, 4, 7 and 9) are involved in normal tissue homeostasis (18–20). Production of selective small molecule inhibitors is complicated by shared domains of the collagenases and the homology of the catalytic site (21). One tested MMP13 “specific” inhibitor PF152 reduced lesion severity in a canine PTOA model (22) but unfortunately caused nephrotoxicity believed to be caused by off-target effects on the human organic anion transporter 3 (which can be circumvented with our proposed RNAi due to sequence specificity and not structural specificity) (23). For these reasons, we hypothesize that selectively targeting MMP13 (which has not been associated with MSS) through delivery of a locally-retained therapy could be an effective and safe approach for blocking the degenerative PTOA process following joint injury.

Local, intraarticular injections are clinically-utilized in OA, yet face unique drug delivery challenges. One of the major barriers is that synovial fluid is continuously exchanged in the joint, causing most drugs to be rapidly cleared into the lymphatic system (24, 25). The synovial vasculature clears small molecules, while the lymphatics drain away macromolecules (26, 27), resulting in joint half-lives ranging 1-4 hours for commonly used steroids (25). These challenges leave an unmet need for OA therapies that are targeted and/or are better retained within the joint after local injection. Targeting of nanoscale particles is one promising approach that has traditionally relied on using chondroitin sulfate, CII-binding peptides, and bisphosphonates (28–30). Chondroitin sulfate and CII-binding peptides target the cartilage matrix to tether/embed the particles and reduce convective transport through synovial fluid flow, while the bisphosphonates are focused on advanced cases of OA and target exposed subchondral bone. Extracellular substrate targeting has been minimally utilized for delivery of small molecules and has not been investigated, to our knowledge, for local retention of biologics with intracellular targets (31).

Here, we sought to target and retain delivery of RNAi therapy against MMP13 to sites of early cartilage damage in OA and to confirm that matrix targeting can provide functional benefit for delivery of intracellular-acting biologics. RNAi is critical in this application because siRNA can be designed to have selective complementarity with MMP13, obviating the enzyme selectivity concerns associated with small molecule inhibitors. The clinical utility of siRNA medicines has been validated by the recent clinical trial success and FDA approval of both Alnylam’s ONPATTRO™ (patisiran) for treatment of hereditary transthyretin-mediated amyloidosis and GIVLAARI™ (givosiran) for acute hepatic porphyria (32, 33). Here, we extended polymeric siRNA nanopolyplexes (siNPs) recently innovated by our research group (34–37) to develop the first targeted form of this carrier that binds to sites of early OA cartilage damage using a collagen type 2 monoclonal antibody (mAbCII). Cartilage CII is more exposed and accessible for binding after injury (38), and the mAbCII antibody has previously been used as a targeted nano-diagnostic for intravitally measuring severity of OA (39). In the current report, we formulated and therapeutically tested mAbCII-targeted siNPs (mAbCII-siNPs) as a locally injectable system that creates an *in situ* depot of MMP13 RNAi nanomedicine in PTOA-afflicted joints. Matrix-targeted delivery of an intracellular-acting biologic such as siRNA represents a significant departure from the convention of targeting internalizing cellular receptors. Herein, we validate the utility of this approach and prove it therapeutically significant as a DMOAD in a model of mechanical PTOA.

## Results and discussion

### Synthesis and conjugation of polymers

The mAbCII-siNPs were synthesized comprising an endosome-escaping, RNA-condensing core and a passivating, colloidally-stabilizing poly (ethylene glycol) (PEG) surface amenable to antibody conjugation (**Figure 1A**). The diblock copolymer that is assembled to generate this system was synthesized through reversible addition-fragmentation chain transfer (RAFT) polymerization of a random copolymer of 50 mol% 2-(dimethylamino)ethyl methacrylate (DMAEMA) and 50 mol% butyl methacrylate (BMA) from a carboxy-PEG-ECT (4-cyano-4 (ethylsulfanylthiocarbonyl) sulfanylpentanoic acid) macro-chain transfer agent (macro-CTA) and verified by NMR (**Figure S1, S2)**. The poly(DMAEMA-co-BMA) (DB) random copolymer block has a balance of hydrophobic BMA and cationic DMAEMA monomers that has been finely tuned to drive NP self-assembly and stabilization (BMA), enable electrostatic siRNA packaging (DMAEMA), and have an appropriate p*K*a and level of hydrophobicity that drives pH-dependent membrane disruptive function in the early endosomal pH range (40–43). The collagen II targeting mAbCII was conjugated to COOH-PEG-ECT by N-hydroxysulfosuccinimide; 1-Ethyl-3-(3- (dimethylamino)propyl) carbodiimide (sNHS/EDC) chemistry. Successful conjugation between PEG-*bl*-DB and mAb-CII to form mAbCII-siNPs was validated by size exclusion chromatography of the conjugate created at a 1:1 molar ratio of antibody to polymer (**Figure 1B**). The resultant polymers were formulated into siNPs by complexation with siRNA at pH 4 followed by raising to physiologic pH (**Figure 1C**). The control groups included bare siNPs and siNPs functionalized with an off-target antibody (mAbCtrl siNPs). “Dual hydrophobization” was also employed in all siNP formulations. This approach combines the hydrophobicity of BMA in the core of the siNP with C16 modification of the siRNA through conjugation to palmitic acid in order to improve siNP stability and gene silencing longevity of action (36, 44).

**Figure 1:**
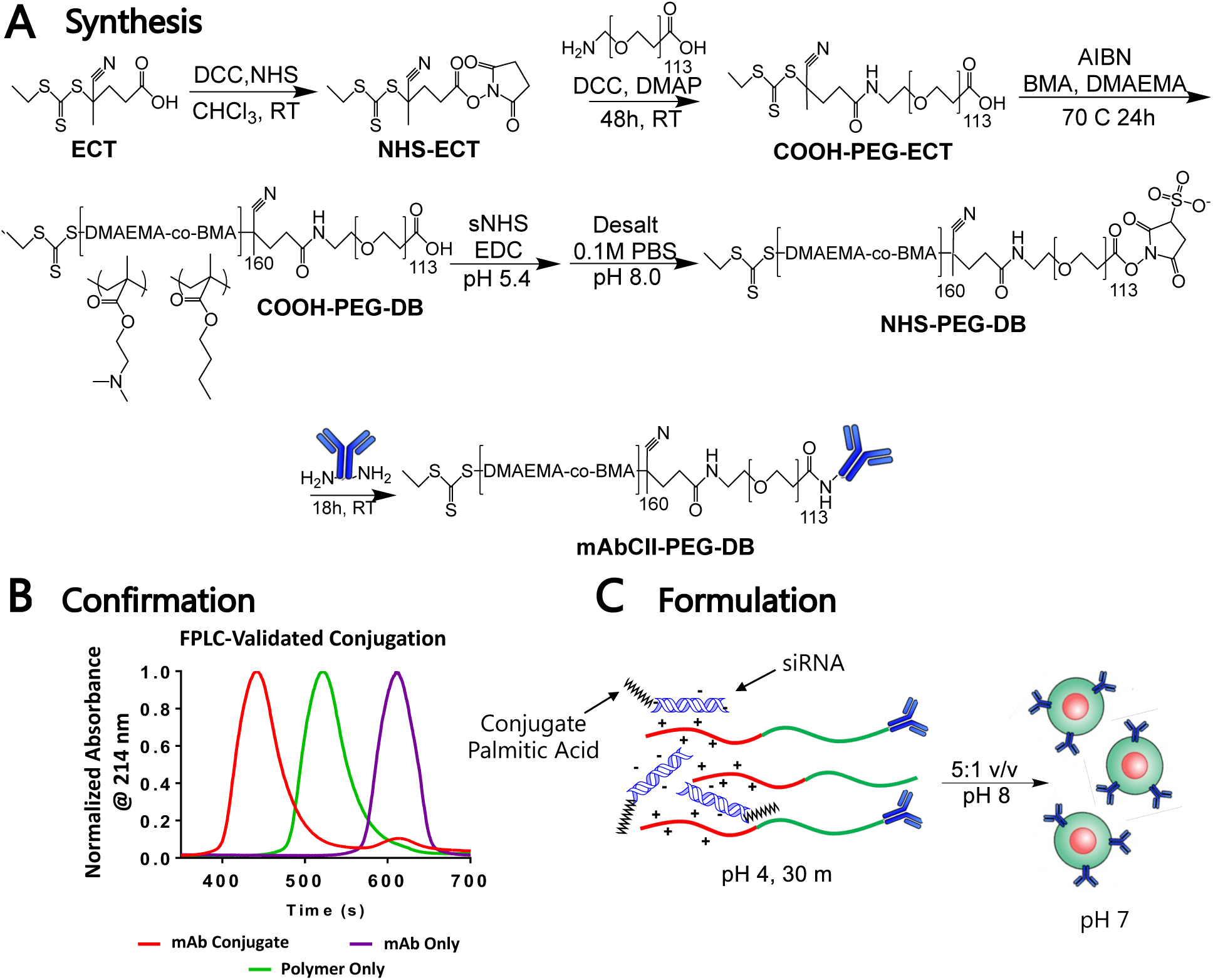
Synthesis of mAbCII-siNPs. (**A**) Polymer synthesis and mAbCII conjugation scheme; (**B**) Size exclusion chromatography confirming mAbCII conjugation to polymer; (**C**) Formulation schematic for siRNA loading and assembly of mAbCII-siNPs.

### Chemicophysical and In Vitro Characterization of mAbCII-siNPs

The siNP hydrodynamic diameter, siRNA encapsulation efficiency, pH-dependent membrane disruptive behavior (as an indirect indicator of endosome escape), and cell viability were assayed for mAbCII-siNPs compared to non-targeted siNPs. The mAbCII-siNPs were statistically equivalent to the non-functionalized siNPs in all these assays (**Figure 2A-D**). The mAbCII-siNPs, prepared at a 1:40 antibody:polymer ratio for optimized targeting (see below), had an average hydrodynamic diameter of 124 nm with a PDI of 1.1 as determined by dynamic light scattering. Encapsulation of siRNA was found to be efficient (∼80%) at N^+^:P^−^ ratios (ratio of positive nitrogen groups in polymer side chains to negative phosphodiester groups in the siRNA backbone) of 10 or above. The hemolysis assay demonstrated significant membrane lysis at the early endosome pH (6.8) and below and negligible activity at extracellular pH (7.4). Cell viability was approximately 80% or greater for doses of 150 nM or less.

**Figure 2:**
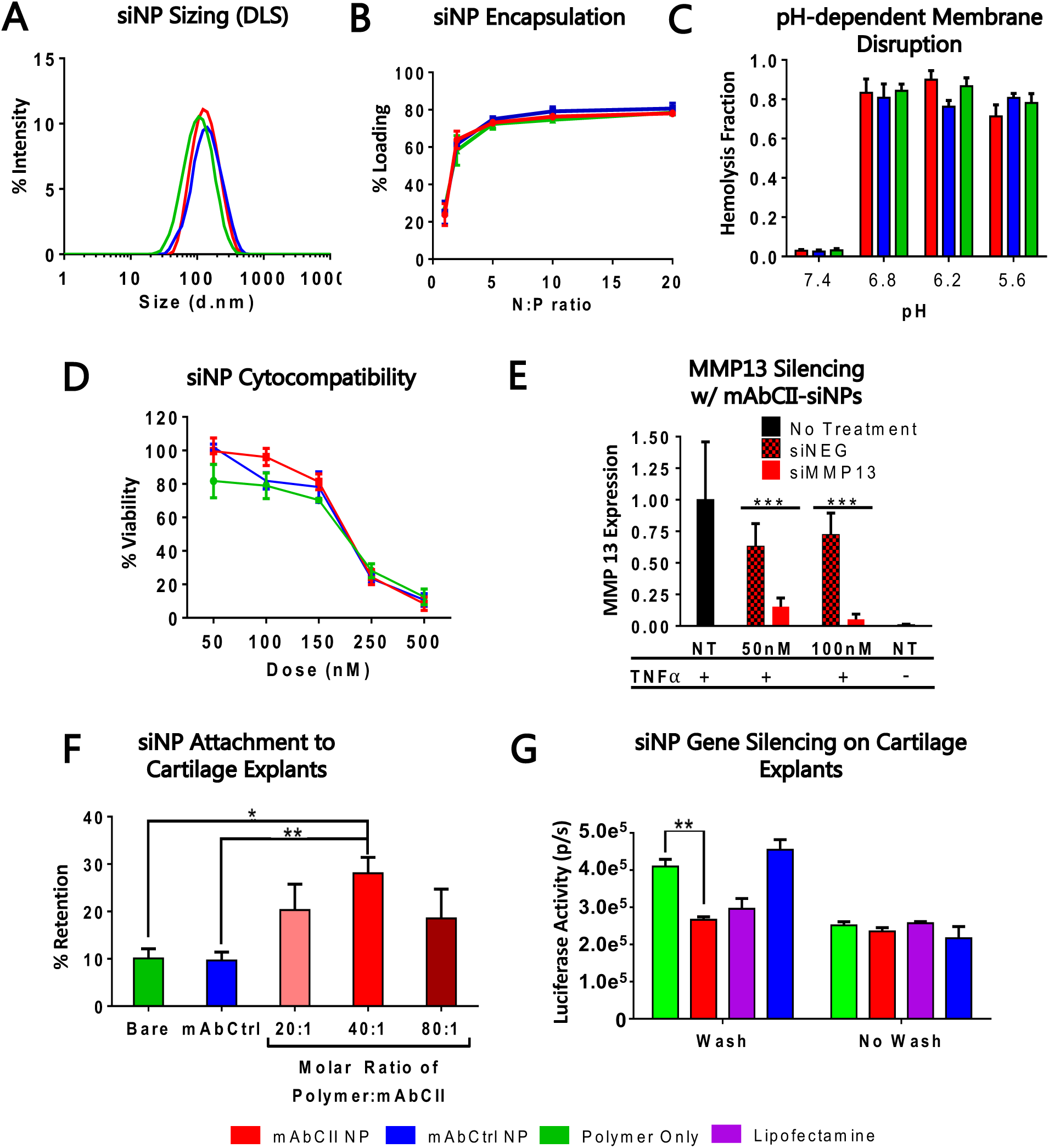
In vitro characterization of siNPs, and comparison of mAbCII-functionalized siNPs to bare and control antibody-functionalized siNPs. (**A**) siNP sizing by dynamic light scattering; (**B**) siRNA encapsulation by Ribogreen assay; (**C**) pH-triggered membrane disruption by hemolysis assay; (**D**) Cytocompatibility of siNPs across a broad dose range; (**E**) MMP13 silencing by lead siRNA candidate in ATDC5 cells stimulated with TNFα using mAbCII-siNPs for delivery; (**F**) Retention of siNPs with varied polymer:mAbCII molar ratios on trypsin damaged porcine cartilage explants after thorough washing; (**G**) Substrate-mediated silencing of MMP13 *in vitro* enhanced by mAbCII-siNP binding and retention on trypsin-damaged porcine cartilage (* = p < 0.05; ** = p < 0.01; *** = p < 0.001).

Silencing of MMP13 was also tested in cultured, chondrogenic ATDC5 cells stimulated with TNFα. The cells were pretreated for 24 h with the siRNA formulations, stimulated with 20 ng mL^−1^ TNFα for 24 h, and then assayed for MMP13 gene expression using TaqMan PCR. Information on screens that identified the leading MMP13 siRNA sequence are found in **Table S1** and **Figure S3**. The best candidate (siMMP13) demonstrated greater than 80% knockdown with a 50 nM dose delivered by the mAbCII-siNPs when compared with a nontargeting siRNA sequence (siNEG) (**Figure 2E**). No substantial change is observed in siNP bioactivity following decoration with a targeting antibody and that the mAbCII-siNPs can achieve potent MMP13 silencing in cells under pro-inflammatory conditions.

### *Ex vivo* CII targeting and substrate-mediated RNAi in ATDC5 cells

The mAbCII-siNPs were assessed for binding to trypsin-damaged porcine cartilage explants. The mAbCII-siNPs were prepared with a range from 20:1 to 80:1 non-conjugated polymer: antibody modified polymer molar ratios. The polymers used comprised rhodamine acrylate (exc/emm: 548/570 nm) copolymerized at 1 mol% in the poly(DMAEMA-*co*-BMA) block to enable fluorescent measurement of carrier retention on the damaged cartilage plugs. Retention of siNPs on trypsin-damaged cartilage after washing with phosphate buffer saline (PBS) was quantified by IVIS imaging and showed that conjugation of mAbCII to the polymer at 40:1 polymer:mAbCII molar ratio provided the best retention performance (**Figure 2F**). These data confirm that mAbCII conjugation enhances binding of siNPs to exposed CII in damaged cartilage and motivated our focus on the 40:1 conjugation ratio for subsequent studies.

Subsequently, the porcine cartilage binding assay was adapted to confirm whether matrix bound mAbCII-siNPs could achieve effective substrate-mediated siRNA delivery and bioactivity. Following incubation of all siNP groups loaded with anti-luciferase siRNA (siLuciferase) with trypsin-damaged cartilage, a PBS washing step was done to simulate synovial fluid clearance. Murine chondrogenic ATDC5 cells that were lentivirally transduced in-house with a constitutive luciferase reporter were then seeded over the damaged cartilage that had been pre-treated with mAbCII-siNPs, bare-siNPs, mAbCtrl-siNPs, or lipofectamine 2000. In parallel, the same groups were run without the washing step to clearly elucidate the benefit of siNP matrix binding and retention. Significantly higher luciferase silencing was observed with mAbCII-siNPs compared to bare and mAbCtrl-siNPs when a wash step was used prior to cell seeding (**Figure 2G**). Following the measurement of luciferase expression, cell viability of each group was determined using the Promega CellTiter-Glo luminescent cell viability assay following Promega’s standard protocol. Only cells treated with lipofectamine 2000 (100 nM) were found to have significantly lower viability than other treatment groups. These data confirm a potential pharmacokinetic benefit of siNP matrix binding and that substrate-mediated delivery of matrix-targeted siNPs achieves target gene silencing.

### *In vivo* targeting-dependent MMP13 silencing in mice

An acute PTOA model of noninvasive repetitive joint loading was used by subjecting the left knee joint of 8-week-old C57BL/6 mice to 50 cycles of compressive mechanical loading at 9N (**Figure 3A**). This procedure was repeated three times per week over a period of two weeks using conditions adapted from previous studies (39, 45). Following loading, mice were treated via intraarticular injection of 0.5 mg/kg per knee of formulated siRNA with mAbCII-siNPs, bare siNPs, or mAbCtrl-siNPs. All forms of siNPs contained a rhodamine acrylate monomer integrated into the poly(DMAEMA-*co*-BMA) block that forms the NP core, enabling IVIS fluorescence imaging to assess pharmacokinetics. Retention in the knees was measured over 72 h, after which knees were excised and the joints were imaged again *ex vivo*. The mAbCII-siNPs had significantly higher retention within the joint compared to mAbCtrl- and bare siNPs (**Figure 3B-D**). It was also supported that the enhanced joint retention was pathology-driven and due to exposed collagen II associated with cartilage damage (46, 47), as retention was also higher in PTOA-damaged knees relative to non-injured knees (**Figure S4**). Finally, in this acute PTOA model, mAbCII-siNP/siMMP13 (candidate sequence from screen; Figure S3) treatment more potently silenced MMP13 expression relative to untargeted siNPs, achieving greater than 90% target gene knockdown in mechanically-loaded PTOA joints 3 days after treatment (**Figure 3E**).

**Figure 3:**
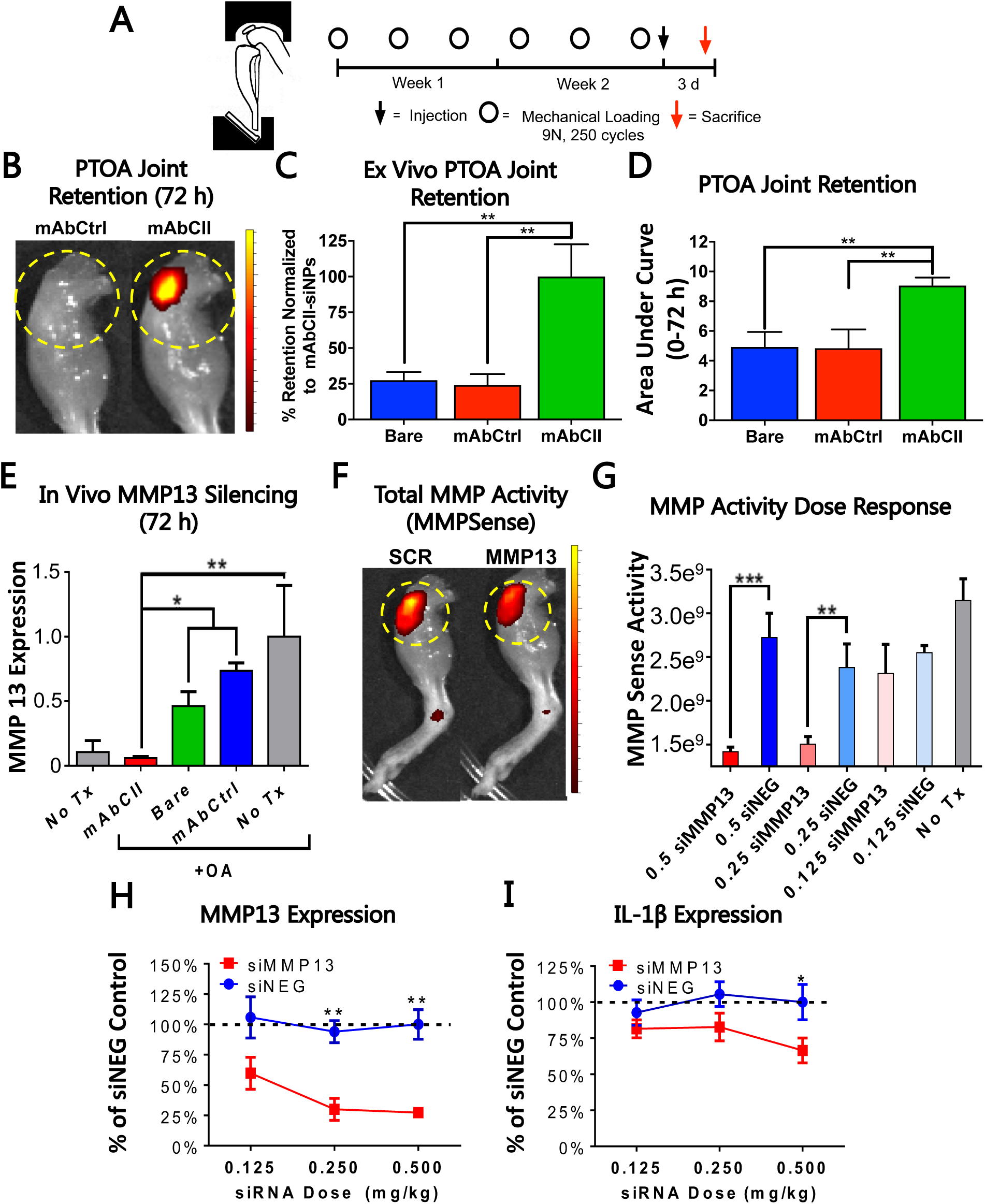
mAbCII-siNPs are better locally-retained in mechanically loaded knees and potently silence MMP13 in short-term in vivo PTOA studies. –(**A**) Schematic illustrations of the mouse knee mechanical loading apparatus and of the loading regimen used in the short-term PTOA model studies; (**B-C**) Representative *ex vivo* imaging and quantification of retention of mAbCII-siNPs compared to mAbCtrl- and Bare siNPs in extracted limbs at 3 days; (**D**) *in vivo* knee retention of mAbCII- and mAbCtrl-siNPs presented as pharmacokinetic area under the curve calculated from intravital imaging over 3 days post-treatment; (**E**) *In vivo* expression of MMP13 measured by TaqMan qPCR in mouse knees treated with 0.5 mg/kg siRNA dose per knee using mAbCII or control (bare and nonbinding control antibody) siNPs; (**F-G**) Example images of total MMP activity as visualized with MMPSense at 0.5 mg/kg siRNA dose and total MMP activity quantified; (**H-I**) Dose dependent effects of mAbCII-siNP delivery of siMMP13 or siNEG (in mg/kg) on expression of MMP13 and IL1β as quantified by TaqMan qPCR (* = p < 0.05; ** = p < 0.01; *** = p < 0.001).

Dose dependent *in vivo* gene silencing activity of mAbCII-siNP/siMMP13 was also measured for siMMP13 (0, 0.125, 0.25, and 0.5 mg/kg per knee injected intraarticularly) or siNEG (0.5 mg/kg) in PTOA knees. Doses were administered after two weeks of cyclic, mechanical loading. At 72 h post-treatment with siNPs, total MMP activity was quantified by IVIS imaging 24 h after intravenous injection with MMPSense (probe activated by MMPs 2, 3, 7, 9, 12, and 13), showing that 0.25 and 0.5 mg/kg siMMP13 doses delivered with mAbCII-siNPs significantly reduced total MMP activity (**Figure 3F-G**). MMP13 and IL-1β expression were quantified in the same experiment by TaqMan qPCR from joint samples collected at 72 h following treatment (**Figure 3H-I**). While both 0.25 and 0.5 mg/kg injections significantly reduced MMP13 gene expression silencing, reduction in IL-1β expression in OA knees was only seen for 0.5 mg/kg, suggesting that this dose provided a broader suppression of the OA inflammatory response. Therefore, a 0.5 mg/kg per knee dose was selected for subsequent, longer-term studies. The mAbCII targeting enhances gene knockdown of MMP13 with roughly 40% greater silencing over bare-siNPs/siMMP13 and 70% greater than mAbCtrl-siNPs/siMMP13, motivating testing in a longer term and more challenging OA model to assess mAbCII-siNPs/siMMP13 therapeutic efficacy as a DMOAD.

### MMP13 silencing in longer-term osteoarthritis mouse model

A 6-week murine study was completed to evaluate the therapeutic effect of weekly doses of MMP13-silencing mAbCII-siNPs in a longer-term and more aggressive PTOA model. Tests for the more aggressive PTOA model were carried out in C56BL/6 mice aged to 6 months and subjected to a more rigorous cyclic mechanical loading protocol of 9N, 500 cycles, 5 times per week, for 6 weeks (**Figure 4A**) (48). Doses of 0.5 mg/kg siRNA were administered into each knee weekly, starting concurrently with mechanical loading. MMPSense and Alexafluor-labeled mAbCII antibody were injected intravenously 24 h before sacrifice to gauge total MMP activity and quantify cartilage damage, respectively. Even though the study takedown and gene expression analysis was done a full week following administration of the final dose, MMP13 expression interference achieved with mAbCII-siNPs was consistent with the short-term model, with greater than 80% reduction compared to OA joints treated with siNEG using mAbCII-siNPs (**Figure 4B**). Immunohistochemical (IHC) staining revealed that MMP13 production both in the articular cartilage and the synovial tissue was reduced by mAbCII-siNP/siMMP13 treatment (**Figure 4C, D**). The inflamed state associated with PTOA is characterized by detrimental tissue restructuring potentiated by cytokines and growth factors produced in the synovium of the joint (49). Degradation products of degraded CII have signaling properties that contribute to catabolic activity, hypertrophy, and apoptosis (50). The mAbCII-based targeting of siMMP13 in the PTOA joint may have both primary, cartilage-protective effects on MMP13 gene silencing and also secondary effects associated with reduced production of degradation products that amplify downstream inflammation and MMP production.

**Figure 4:**
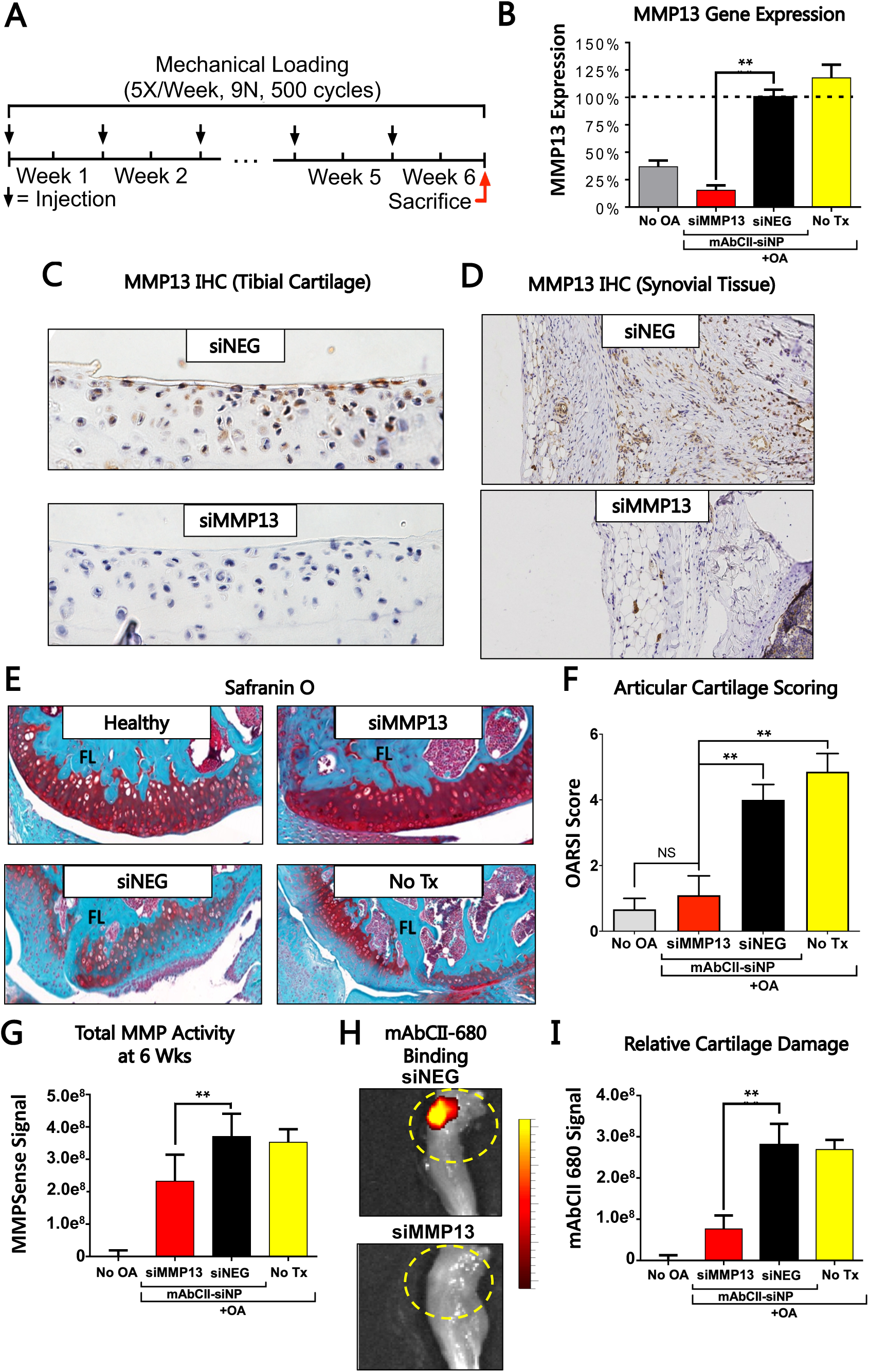
Long-term MMP13 silencing reduces MMP13 protein levels in cartilage and synovium and protects mechanically-loaded joints from OA progression. (**A**) Loading and treatment regimen used in the long-term mouse model; (**B**) MMP13 expression at the end of week 6; (**C-D**) IHC for MMP13 showing that targeted siMMP13 treatment reduced MMP13 protein levels both in the articular surface of the tibia and in the synovial tissue; (**E-F**) Representative Safranin O staining of the articular surface of the femur and quantification of cartilage damage by the OARSI osteoarthritis cartilage histopathology assessment system. (**G**) Total MMP activity at 6 weeks measured by MMPSense. (**H-I**) Representative images and quantification of the binding of fluorescently labeled mAbCII in the joint at 6 weeks as a marker for relative cartilage damage and resultant superficial exposure of Collagen II (* = p < 0.05; ** = p < 0.01; *** = p < 0.001).

Histological analysis showed that mAbCII-siNP/siMMP13 treatment significantly reduced PTOA-associated joint structural changes. Coronal sections of fixed knee joints were stained with Safranin O and Fast Green to evaluate cartilage histopathology. Sections were then blindly scored by a trained pathologist (**Figure 4E-F**) using criteria detailed in the supplement (**Table S2A, S2B**). Safranin O stains proteoglycans associated with normal cartilage a deep red and is used for histopathological scoring of cartilage as recommended by the Osteoarthritis Research Society International (OARSI) (51). The reduced Safranin O staining and surface discontinuity observed in untreated OA and siNEG-treated OA is indicative of proteoglycan loss and cartilage erosion and was significantly prevented in siMMP13-treated joints.

Total MMP activity was also significantly reduced in the mAbCII-siNP treated animals compared to controls that were either untreated or treated with mAbCII-siNPs loaded with siNEG (**Figure 4G**). Relative binding of Alexafluor-labeled mAbCII antibody to pathologically exposed CII is a biomarker for degree of cartilage damage (39, 52, 53) and was thus used here as an intravital readout. Significantly greater binding of mAbCII was observed in mAbCII-siNP/siNEG treated and untreated mice compared to mAbCII-siNP/siMMP13 treated mice, indicating that targeted MMP13 silencing protected the cartilage architecture (**Figure 4H, I**). Levels of Alexafluor680-mAbCII binding in the PTOA joint were similar between untreated mice and mAb-CII-siNP/siNEG treated mice, indicating that treatment with mAbCII-functionalized siNPs one week prior did not in itself interfere with the subsequent mAbCII-based cartilage damage measurement.

Hematoxylin and eosin (H&E) staining was used to evaluate overall joint status, including synovial response (**Figure 5A**). The loading protocol utilized induced robust synovial thickening and ectopic mineralization. Whole joint histology was blindly scored by a pathologist (**Figure 5B**) based on the degenerative joint disease criteria outlined in the supplemental information (**Table S2**). While cartilage structure was fully protected by mAbCII-siNP/siMMP13 treatment (treated mice had statistically equivalent OARSI score to normal mice with no load-induced PTOA), there was also a significant protection of other joint tissues, including less synovial hyperplasia compared to control-treated animals. The H&E joint sections for siNEG and untreated groups also appeared to contain higher ectopic calcification in the synovial membrane compared to the mAbCII-siNP/siMMP13 group. To more globally visualize and quantitatively assess ectopic calcification of the synovium, we utilized microCT, revealing that mAbCII-siNP/siMMP13 treatment protected against synovial mineralization (**Figure 5C-D**). These data support that MMP13 silencing does have secondary benefits on the joint beyond directly reducing articular cartilage loss.

**Figure 5:**
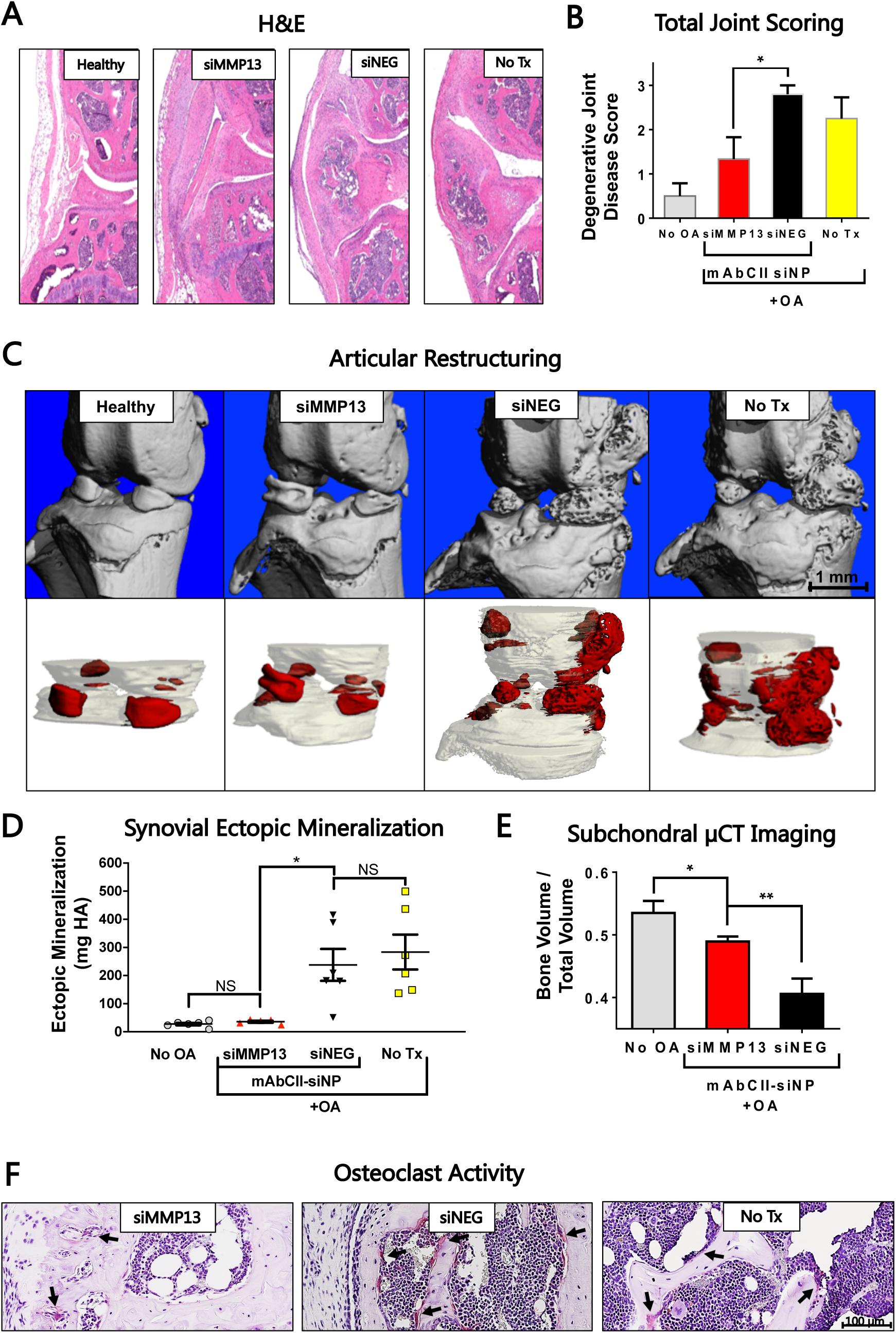
mAbCII-siNP/siMMP13 silencing decreases joint destruction by reducing synovial response and subchondral bone loss. (**A-B**) H&E staining of joints, focusing on the synovial regions, and average Degenerative Joint Disease Score by treatment group. Blinded histological scoring was completed by a trained histopathologist; (**C**) 3-dimensional renderings of synovial ossification in the joint space from microCT scans; (**D**) Quantified degree of ectopic synovial ossification in quantified hydroxyapatite (HA); (**E**) microCT-derived bone volume fraction of the femoral and tibial subchondral bone; (**F**) TRAP staining of PTOA groups highlighting osteoclast activity in the subchondral bone (Arrows highlight active osteoclasts; * = p < 0.05; ** = p < 0.01; *** = p < 0.001).

When evaluating OA joints clinically, the presence of osteophytes and ossified nodules within the synovium is especially important, as these characteristics are used to gauge OA progression and appear in the most advanced stages of the disease (54). These rigid calcium deposits are associated with synovial macrophage activation in experimental OA (55) where they concentrate local mechanical stress. Further, presence of calcium phosphate can exacerbate synovitis by activating inflammasomes and consequently triggering production of the OA driver IL-1β (56). While inflammation affects patient comfort, complications and acute pain from osteophytes and calcium deposits are often cited as primary reasons for advanced OA patients to cease exercise altogether and to resort to total knee replacement (57). Inhibiting pathological ossification would be anticipated to enable maintenance of an active lifestyle and to delay total knee replacement.

To look more broadly at the joint and surrounding tissues, subchondral trabecular bone was also characterized using microCT. Interestingly, mAbCII-siNP/siMMP13 treatment significantly reduced loss of subchondral trabecular bone volume associated with PTOA (**Figure 5E**). TRAP (tartrate-resistant acid phosphatase) staining was also performed to assess osteoclast activity as a snapshot of bone resorption (**Figure 5F**). Activated osteoclasts were increased in untreated, mechanically loaded knees compared to those receiving mAbCII-siNP/siMMP13 treatment. These data reinforce the complexity of OA as a full joint disease, while also indicating that MMP13 RNAi has beneficial, global impacts within the joint by also impacting cartilage crosstalk with surrounding tissue. There is strong clinical precedent for observing multiple, concomitant types of aberrant mineral homeostasis within the OA joint. Clinically, calcium “crystals” often manifest in the synovial tissue and fluid. At the same time, pathological vascularization and thickening of the subchondral bone plate can cause loss of subchondral trabecular bone (56). Animal models and human samples suggest an influence of MMP13 and cathepsin K, specifically, in driving pathological subchondral bone resorption in late-stage OA in human tissues (58). In accordance with these studies, we find that subchondral bone resorption and pathologic bone restructuring within the whole joint are reduced with MMP13 inhibition.

### Benchmark comparisons to clinical gold standard steroid treatment

The proposed therapy is positioned as an early intervention for PTOA. The most prevalent clinical intervention beyond the use of oral NSAIDS is the intraarticular administration of steroids, which are recommended to be given up to four times per year (59). Of corticosteroids, the most commonly used is methylprednisolone (60). Because the longer-term, 6-week osteoarthritis mouse model used in our therapeutic studies is aggressive, benchmarking was compared against both a single dose of methylprednisolone at the time of first injury (most similar to frequency of dosing used clinically) and a weekly dose, with the latter done to match the protocol used for testing of mAbCII-siNPs/siMMP13 (dosing described in Supplemental Information).

Histological sections on joints extracted at the endpoint of this study were blindly scored by a pathologist following tissue staining with hemotoxalin/eosin and safranin-O (**Figure 6A-D**). Neither single or weekly methylprednisolone injections significantly altered either the OARSI or Degenerative Joint Disease Scores relative to untreated PTOA knees. These findings are in agreement with other studies demonstrating that while steroids temporarily alleviate inflammatory pain, there is no demonstrated protection of cartilage structural integrity (14). Notably, mAbCII-siNPs/siMMP13 treatment significantly reduced both the DJD and the OARSI score relative to controls, while neither single or weekly steroid treatment provided any significant protection by these joint scoring metrics. Similar to the histological scoring outcomes, neither steroid treatment protocol blocked ectopic mineralization in the synovium, unlike mAbCII-siNP/siMMP13 therapy (**Figure 6E**). These distinctions further highlight the therapeutic potential of specific inhibition of MMP13, a molecular driver that broadly underlies several aspects of PTOA joint destruction.

**Figure 6:**
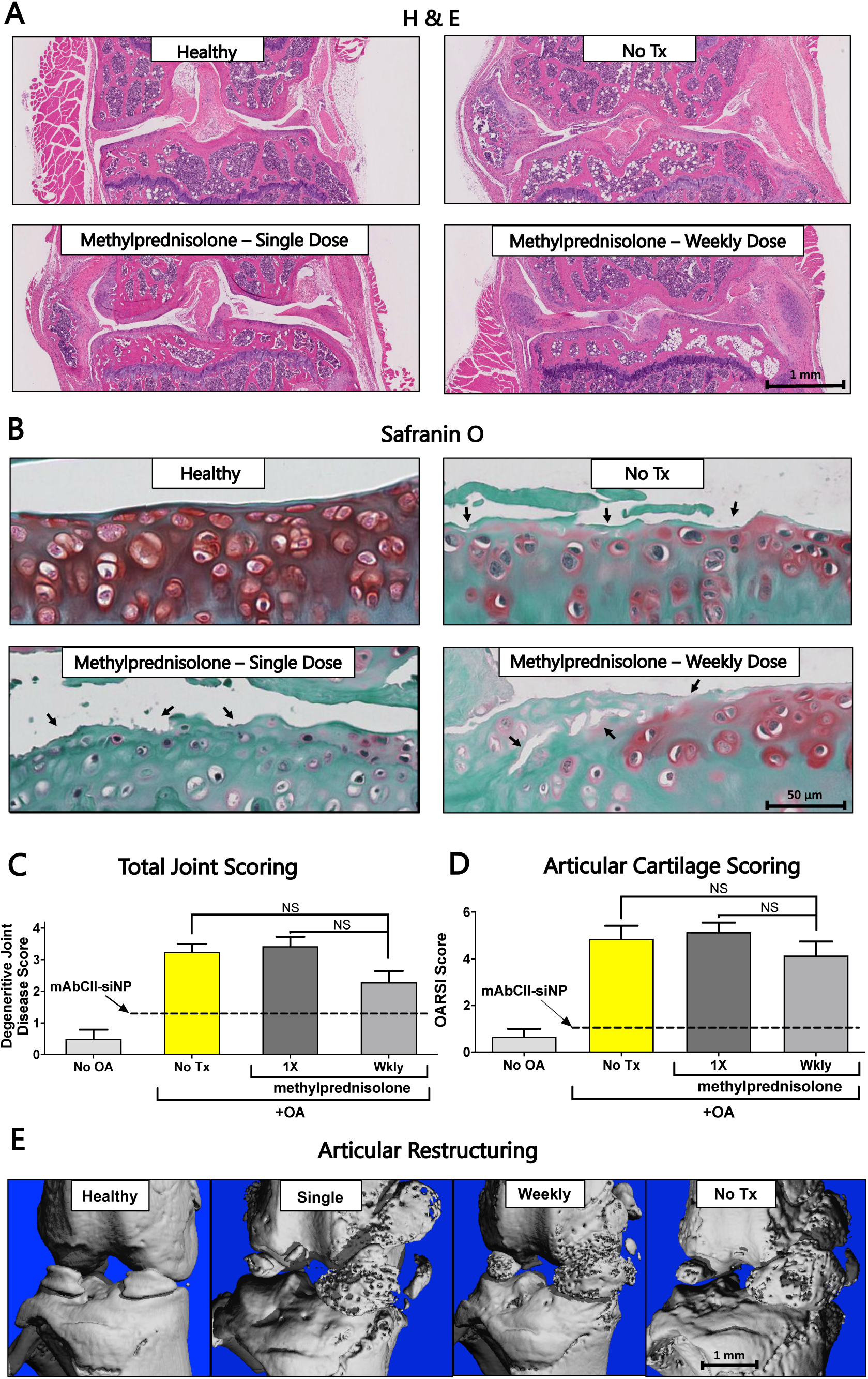
Steroid treatment does not provide DMOAD effects in relation to mAbCII-siNP/siMMP13. (**A**) Representative whole joint histological sections stained with H&E; (**B**) Representative safranin-O staining of the articular surface of the tibia and femur; (**C**) Total joint scores were determined from H&E stained slides by a treatment-blinded histopathologist; (**D**) Cartilage damage by the OARSI osteoarthritis cartilage histopathology assessment system was also determined by a treatment-blinded histopathologist; Arrows highlight focal cartilage damage and regions of cartilage degeneration; (**E**) MicroCT imaging of ectopic mineralization in the synovial tissues; * = p < 0.05; ** = p < 0.01; *** = p < 0.001).

### MMP13 silencing broadly affects expression of OA-associated genes

Targeting MMP13 directly by RNAi successfully silenced MMP13 and was found to have broader effects on overall joint health, including in the synovium and subchondral bone. To further characterize the global impacts of targeted MMP13 inhibition, the extended mechanical loading mouse model was repeated with the same parameters except that samples were harvested at 4 rather than 6 weeks to capture a more intermediate stage of disease (**Figure 7A**). The nanoString nCounter Inflammation panel was used to quantify expression of 254 genes in the knee joint samples (articular cartilage and synovial tissue combined at a consistent 1:1 mass ratio). Unsupervised analysis using nanoString software indicate that the joints treated with mAbCII-siNPs/siMMP13 were closer to normal tissue and more different from untreated OA tissue than mAbCII-siNPs/siNEG and more (**Figure 7B**). Compared to treatment with mAbCII-siNPs/siNEG, treatment with mAbCII-siNPs/siMMP13 significantly suppressed expression levels of several clusters of genes with notable associations with OA progression (**Figure 7C,D**).

**Figure 7:**
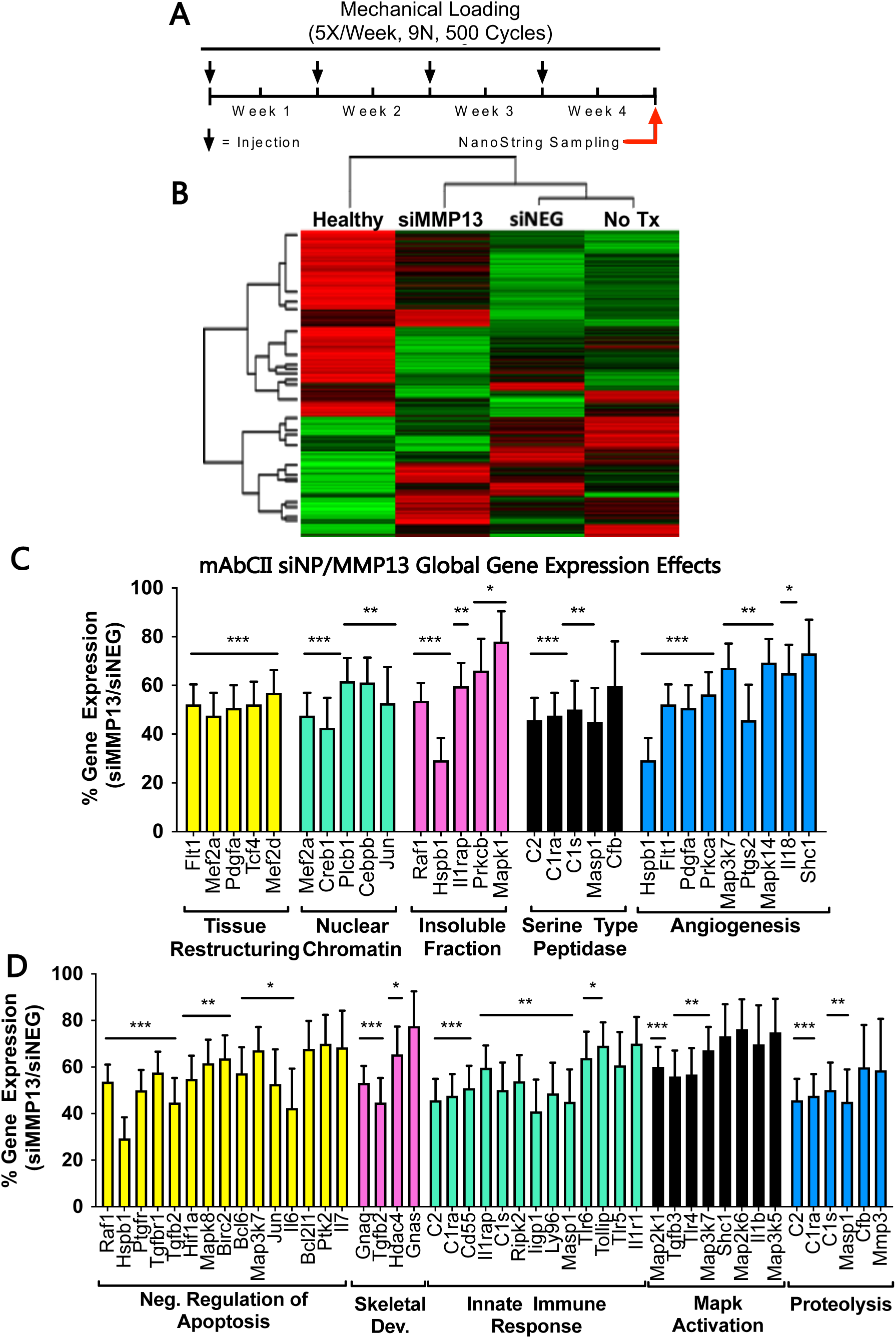
MMP13 silencing globally impacts gene expression patterns within mechanically-loaded PTOA joints. **A**) Loading and treatment regimen for the nanoString analysis study; (**B**) Unsupervised sorting of treatment groups from most to least similar to normal joints as quantified by nanoString at the end of week 4. Gene expression is shown as high- (green) or low-expression (red) sorted vertically by differences between treatment groups; (**C-D**) Gene clusters were significantly different between siNEG- and siMMP13-treated joints (5 most changed in **C**, next 5 most changed in **D**). The plots are arranged such that the clusters’ p-values between siMMP13 and siNEG treatment groups is lowest on left and highest on right (* = p < 0.05; ** = p < 0.01; *** = p < 0.001).

In PTOA joints, mAbCII-siNPs/siMMP13 treatment significantly reduced clusters of genes associated with tissue restructuring and angiogenesis. Tissue remodeling is an active process in response to injury, in the articular cartilage and especially in the synovium where capsule thickening, vascularization, and hyperplasia occurs. These processes are associated with the upregulation of genes such as MEF2α (chondrocyte hypertrophy) and PDGFα (a potent synovial fibroblast growth factor) (61, 62), which were suppressed by mAbCII-siNPs/siMMP13. MMP13 silencing also suppressed Flt-1, which is a VEGF receptor, predominantly expressed by vascular endothelial cells and involved in angiogenesis. The more active tissue remodeling and thickening processes in control PTOA joints was also indicated by mAbCII-siNP/siMMP13 treatment-associated suppression of gene clusters related to proteolysis and skeletal development. MMP13 silencing also led to downregulation of genes associated with apoptosis, such as Jun (which also induces MMP13) and the inflammatory cytokine IL6, which is mechanistically involved in propagation of cellular stress and synovial inflammation (63, 64).

Treatment-associated downregulation of genes related to tissue restructuring, apoptosis, angiogenesis, and proteolysis were also complemented by reduction of innate immune activation in joints with targeted MMP13 silencing. OA-relevant PTGS2 (prostaglandin-endoperoxide synthase 2, an inflammation driver that encodes the cyclooxygenase 2 enzyme (COX2) and IL1RN encoding the interleukin 1 receptor antagonist protein (IL1RaP) in which its knock-out correlates with spontaneous arthritis development (65) were, for example, significantly reduced in mechanically loaded joints with mAbCII-siNP delivery of siMMP13 versus siNEG. Coupled with reduction of innate immune response, MMP13 silencing also reduced expression of several serine proteases (C1S, C1RA, and C2) that are components of the complement pathway, which is known to be associated with OA progression (66, 67). The local reduction in these components indicates reduced number or activation of circulating monocytes, macrophages, and monocyte-derived dendritic cells that all express C1s, C1RA, and C2 (68). MMP13 silencing also reduced expression of IL-1β (MAPK activation cluster), an inflammatory cytokine that drives OA driver by inducing nitric oxide production, increasing synthesis of MMPs and aggrecanases, while suppressing proteoglycan synthesis (69). Interestingly, recent work has highlighted how macrophages embedded in synovial tissue serve to induce a proinflammatory state (70) and, in combination with synovial fibroblasts, appear to be the lead contributors to MMP13 expression within the joint space as observed by MMP13 IHC. Not only does MMP13 inhibition directly reduce cartilage degradation, it has secondary impacts on overall joint health by disrupting the “degenerative cycle” that is perpetuated by degradative products of the cartilage matrix (17).

The global changes to inflammatory gene expression observed in this study, in combination with joint analysis utilizing microCT and histology, all support the therapeutic value and safety of local, MMP13 silencing targeted to cartilage matrix. It is thought that MMP13 activity within the synovial joint drives OA progression (71). By contrast, MMP13 expression by osteocytes is important for normal perilacunar/canalicular remodeling (PLR). Selective genetic ablation of MMP13 expression in osteocytes disrupts subchondral bone homeostasis in a way that exacerbates subchondral bone sclerosis and OA pathology (72). We speculate that a CII-targeted delivery system such as mAbCII-siNPs that is injected directly into the intraarticular space will localize effects to cartilage and possibly adjacent synovial tissues, while any unbound treatment would clear primarily through lymphatic drainage (73), limiting exposure and gene targeting effects on cells embedded within the more remote subchondral bone tissue shielded by mineralized matrices. The concept that mAbCII-siNPs/siMMP13 helped to maintain normal mineralization homeostasis within the joint is supported by the microCT analyses showing that treated animals maintained a more normal subchondral bone volume fraction with reduced abnormal ectopic mineralization versus control treatment. Our matrix-targeted mAbCII-siNPs enable a desirable combination of safe and effective therapeutic MMP13 targeting that may be difficult to achieve with systemic delivery or through delivery of small molecule inhibitors that more readily diffuse into tissues outside of the synovial joint.

### Conclusions

RNAi silencing of MMP13 using matrix-targeted nanocarriers to prolong retention within the osteoarthritic joint provides significant therapeutic benefit in blocking PTOA progression. This study validates the unique concept that matrix targeting for local retention of an in situ formed nanoparticle-based depot is a viable strategy to improve potency and longevity of action of intracellular biologics such as siRNAs. While formulation of larger sized (micro-scale) particles may also facilitate retention, it would not be anticipated to be effective for intracellular acting drugs because of endocytosis, endosome escape, and tissue penetration limitations of larger particles (74). The current system, which uses antibody targeting to reduce joint clearance, achieved as high as 90% target gene silencing in vivo with gene silencing remaining potent even 1 week after the final treatment in a 6-week study. Furthermore, local retention of the injected dose and specific targeting of MMP13 are anticipated to reduce the toxicity concerns that have become associated with systemically-delivered, non-selective synthetic small molecule MMP inhibitors. The therapeutic relevance of this approach is exemplified in an aggressive, long-term PTOA model which showed a treatment-associated 80% reduction in MMP13 expression (p = 0.00231), protection of cartilage integrity (0.45+/−.3 vs 1.6+/−.5 on the OARSI scale; p = 0.0166), improvement in synovial histopathology (1.3+/−.6 vs 2.8+/−.2 on the Degenerative Joint Disease scale; p < 0.05), and less destruction of subchondral bone (16% greater bone volume over total volume; p < 0.01). There were also global gene expression changes associated with MMP13 silencing that suggests this treatment produces both primary benefits through cartilage protection and secondary benefits from blocking inflammatory, degradative feedback loops activated by the release of cartilage degradation by-products. In sum, matrix-targeted delivery for potent MMP13 gene silencing is a promising experimental DMOAD that warrants additional preclinical development because of its significant advantages over current treatments.

### Experimental section

Experimental details and characterization of all particles, animal procedures, and therapeutic assessment are included in Supporting Information.

## Supporting information

Experimental methods are available in the supplementary document.

## Acknowledgements

The authors acknowledge the assistance of the Vanderbilt Translational Pathology Shared Resource (TPSR). TPSR is supported by NCI/NIH Cancer Center Support Grant 2P30 CA068485-14. Dynamic light scattering was conducted at the Vanderbilt Institute of Nanoscale Sciences and Engineering. Bone analysis by microCT was supported in part by the NIH (S10RR027631-01). The technical assistance of Carrie B. Wiese, Joshua R Johnson, and Ray Mernaugh is also acknowledged.

We are grateful to the DOD (DOD CDMRP OR130302), NIH (NIH R01 CA224241, NIH R01 EB019409), NIH (NIGMS T32GM007347), the VA Merit Award BX004151, and the National Science Foundation Graduate Research Fellowship Program (NSF GRF #2016212929) for support.

## Experimental section

### Materials

Unless otherwise stated, materials and reagents were purchased from Fisher Scientific (Waltham, MA, USA) or Sigma-Aldrich (St. Louis, MO, USA).

### Synthesis and conjugation of polymers

*N*-hydroxysuccinimide-functionalized 4-cyano-4 (ethylsulfanylthiocarbonyl) sulfanylpentanoic acid (NHS-ECT) synthesis was verified by NMR (**Figure S1**), and the product was then conjugated to an amine-carboxy bifunctional 5kD PEG to form carboxy-PEG-conjugated ECT for use as an initial chain transfer agent for RAFT polymerization (40). A co-polymer of DMAEMA (2-(Dimethylamino)ethyl methacrylate) and BMA (butyl methacrylate) was chain extended from the COOH-PEG-ECT with a desired target degree of polymerization of 150 (1:1 molar ratio DMAEMA:BMA) to create PEG-DB which was verified by NMR (**Figure S2**). The reaction was purged with nitrogen for 30 minutes. AIBN was utilized as an initiator (10:1 CTA:Initiator ratio) in 10% w/v dioxane. The reaction was stirred at 65 °C for 24 h before precipitation into ether and vacuum drying for 24 h. Polymer was then dissolved and dialyzed in methanol for 48 hours before transition to dialysis in water for another 48 hours.

**Figure S1.**
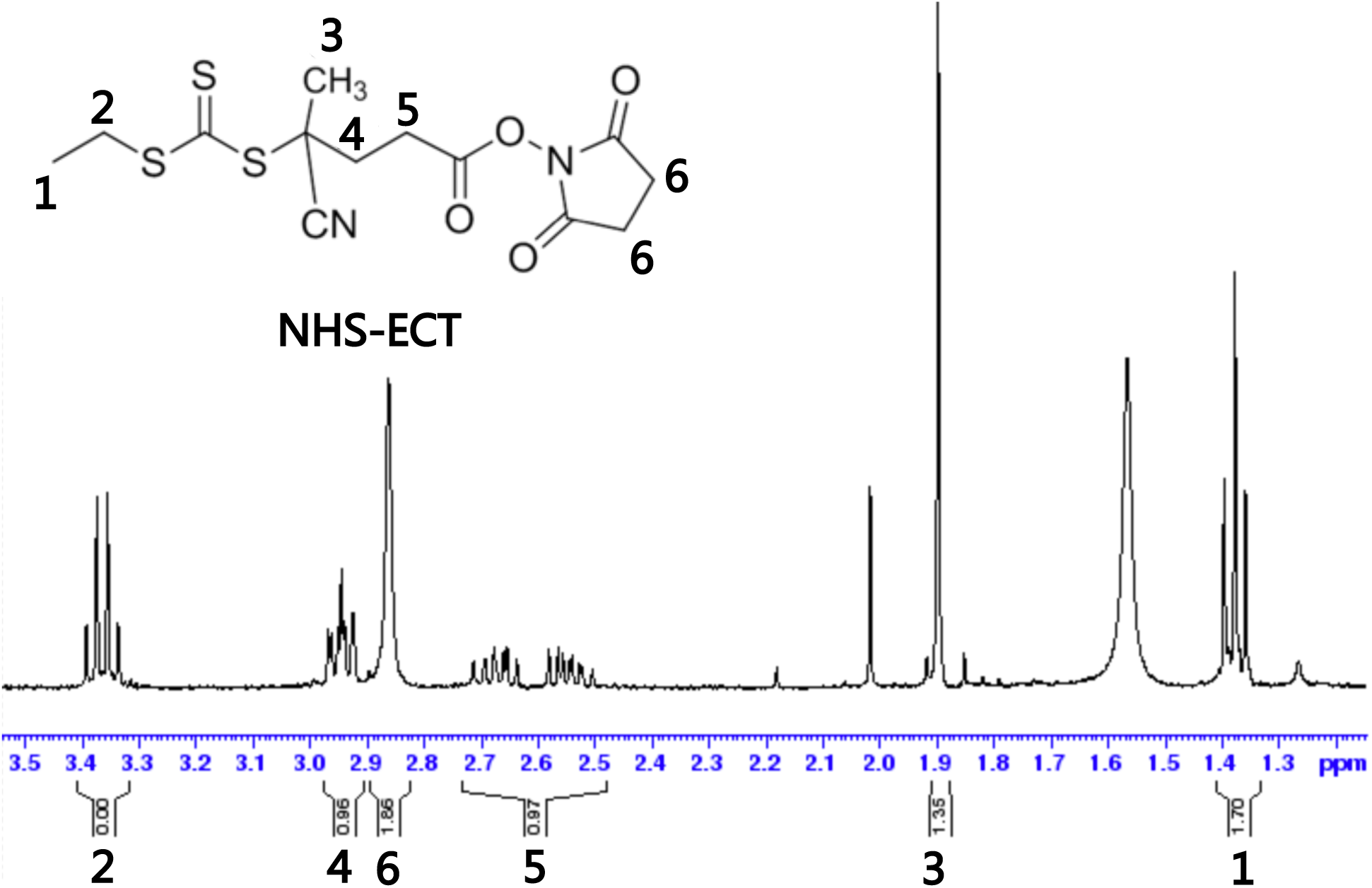
^1^H-NMR spectrum of NHS-ECT, a functionalized RAFT chain transfer agent in CDCl3

**Figure S2.**
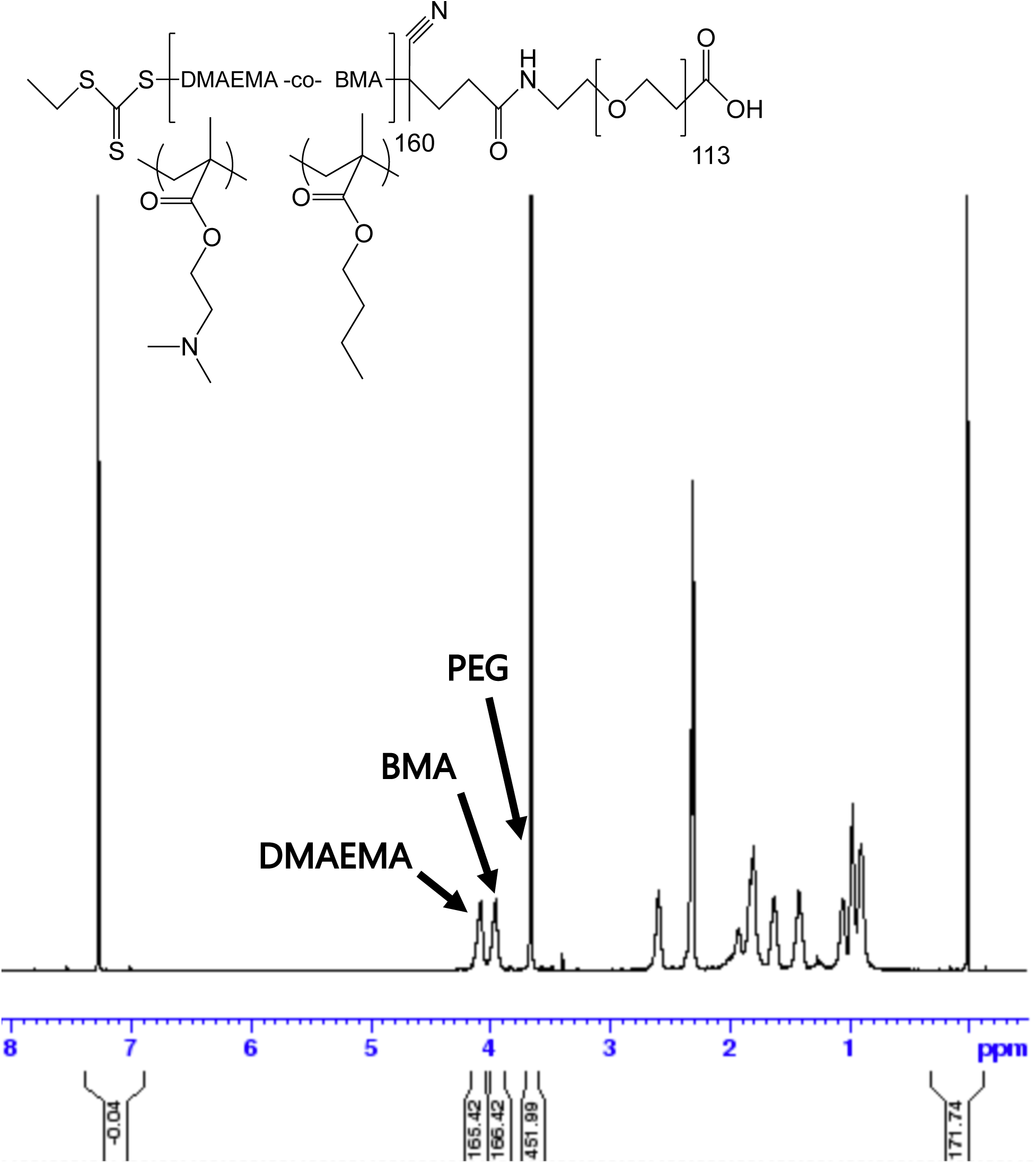
^1^H-NMR spectrum of DB-PEG-COOH in CDCl3 calibrated to the PEG proton content

Subsequently, a two-stage sNHS/EDC conjugation protocol was optimized for activating the polymer and removing excess activating compounds before adding the antibody to avoid antibody crosslinking. Carboxyl terminated PEG-DB polymer was dissolved in ethanol at a 20 mg/mL concentration before addition to 0.05 M MES buffer, pH=6.0 to prepare a final 1 mg/mL solution of COOH-PEG-DB polymer. EDC and sNHS were added at 250 and 500 mM and allowed to react for 15 minutes at room temperature. Excess EDC and sNHS were then eliminated using 10kD MWCO spin filters from Amicon, centrifuging at 3,000 rcf for 13 minutes from an initial volume of 6 mL. Volume lost was replaced with more MES buffer reestablishing initial volume. Antibody at 1 mg/mL in 0.1M PBS, pH=8.0 was added to the activated polymer solution and allowed to react for 18 h at room temperature. Conjugation was verified by size exclusion chromatography as shown above, and fully conjugated mABCII-PEG-DB were mixed with non-functionalized PEG at the appropriate ratio (usually 1:40 conjugated to nonfunctionalized polymer).

### Characterization of particles

Antibody conjugation to polymer was verified by size exclusion chromatography, tracking polymer elution by absorbance at 214 nm through Enrich SEC 650 columns at a flow rate of 0.25 mL/min in 10 mM PBS at pH 8.

A sequence of tests was then conducted to verify that the particle functionality was uncompromised by conjugation. The efficiency of siRNA encapsulation N^+^:P^−^ ratios was evaluated using a Quant-iT Ribogreen assay kit (ThermoFisher Scientific, Waltham, MA). siNP size was evaluated using dynamic light scattering (DLS) (Zetasizer Nano ZS, Malvern, USA). Polymer pH-dependent membrane disruptive function as a marker for endosome disruption/escape was evaluated using a hemolysis assay, as described previously (75).

### Selection of MMP13 siRNA

Seven candidate siRNA sequences targeting different sites of the MMP13 gene were first screened in ATDC5s stimulated with the inflammatory cytokine TNFα (20 ng/mL). Oligonucleotides used in these studies were purchased from Integrated DNA Technologies (Coralville, IA, USA) or Dharmacon, Incorporated (Lafayette, CO, USA). The selected sequence was synthesized with 2’ *O*-methyl modification for enhanced *in vivo* activity.

**Table S1.**
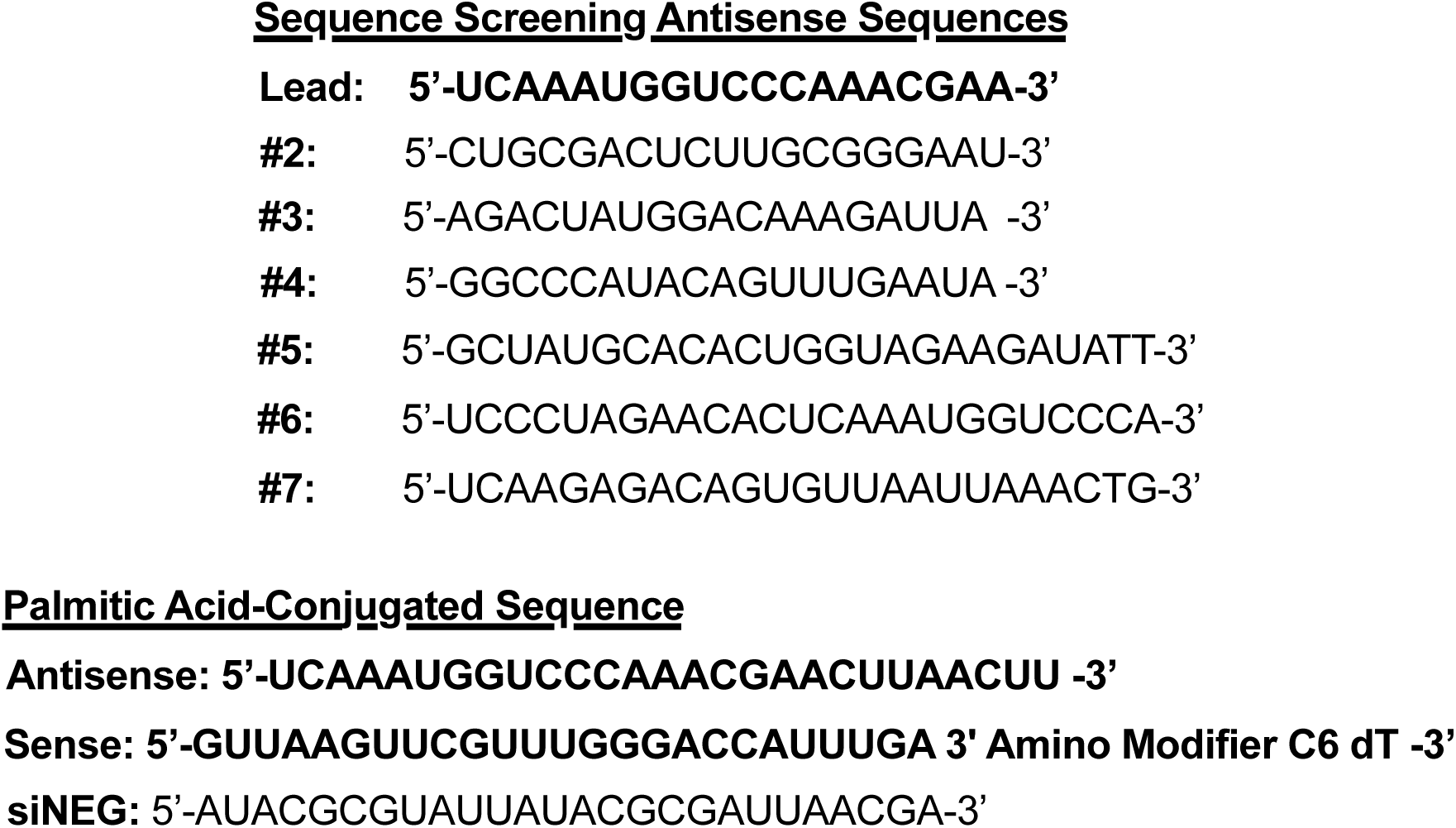
Candidate sequences screened for MMP13 knockdown in inflamed ATDC5 cells. Selected sequence is bolded. The palmitic acid-conjugate modification employed thereafter is also described.

**Figure S3.**
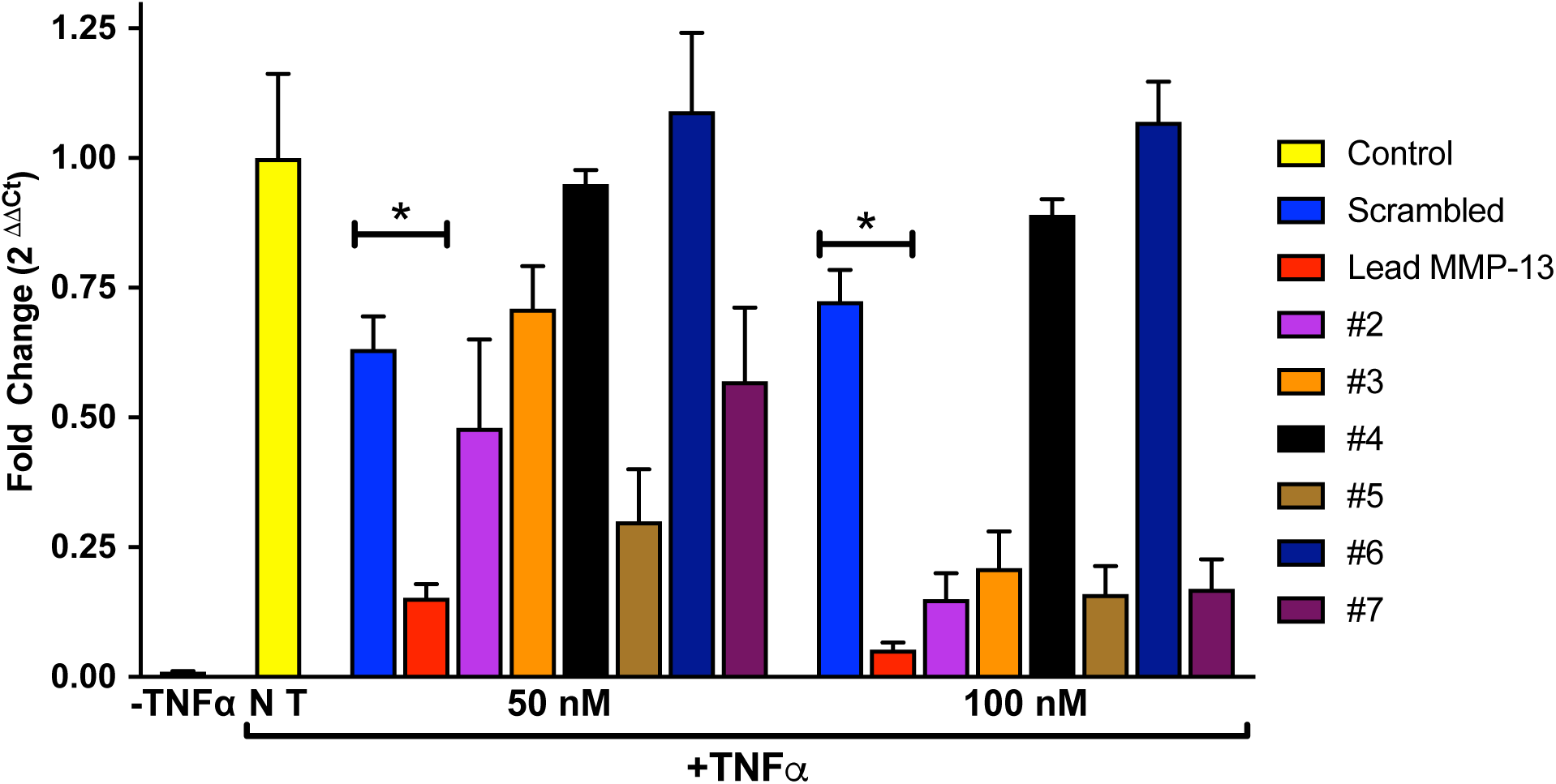
Candidate sequences screened for MMP13 silencing in TNFα-stimulated ATDC5 cells (* = p < 0.05; ** = p < 0.01; *** = p < 0.001).

### Formation of mAbCII-siNPs

Polyplexes were formed by dissolving polymers in 10 mM citric acid buffer (pH 4) before complexation with siRNA for 30 minutes. Polymer was initially dissolved at 3 mg/mL concentration. The siRNA (or palmitic acid-modified siRNA) were complexed at the N:P ratio of 20. Following complexation, the pH was neutralized to 7.4 with sodium phosphate buffer (10 mM; pH 8; 5:1 v/v ratio).

For *in vivo* experiments, siNPs were formed under the same conditions and concentrated using 50kD MWCO 15 mL Amicon spin filters, washing with PBS, and sterile-filtered before injection.

### Cell culture

ATDC5 cells were cultured in DMEM / F-12, GlutaMAX medium with 10% FBS, 1% penicillin/streptomycin at 37 °C in 5% carbon dioxide. Relevant experiments were performed at 80% confluency.

### Cell viability studies

Cytotoxicity was performed using the CellTiter-Glo assay, in accordance with the manufacturer’s protocol.

### Luciferase gene silencing assay

For *in vitro* luciferase knockdown assays, ATDC5 cells were lentivirally transduced with constitutively-expressed luciferase gene in a manner previously described (37). Cells were then seeded at 2,000 per well in 96-well black plates, clear-bottom. After allowing cells to adhere for 24 hours, siNPs were then introduced into cell media at a concentration of 100 nM siRNA (siLUC or siNEG). Treatments were removed after 24 hours of incubation, and cell bioluminescence was then measured on an IVIS Lumina III imaging system (Caliper Life Sciences, Hopkinton, MA) at 24 and 48 hours after treatment by addition of 150 μg/mL luciferin. Luminescence was normalized to that of siNEG NP controls. Finally, cell viability was measured by comparing luminescence of siNEG controls to untreated cells.

### Quantitative reverse transcription PCR

Real-time qRT-PCR was performed utilizing TaqMan primers and reagents, and conducted under conditions outlined by the manufacturer. (Thermofisher Scientific, Waltham, MA; GAPDH: Mm99999915_g1, ACTB: Mm02619580_g1, MMP13: Mm00439491_m1, IL6: Mm00446190_m1)

### In vitro collagen-targeting, uptake, and viability in ATDC5 cells

In order to test targeting and potency of the mAbCII-siNPs formulation, a reverse transfection assay was employed. First, damaged articular cartilage “model lesions” were created by partial trypsin damage of porcine cartilage with 2.5% trypsin for 15 min at 37 °C. Then, each plug was inserted into a well plate and treated with mAbCIIsiNPs or controls. Damaged tissues were incubated for 1 h with mAbCII/siLuc or control siNPs complexed with luciferase silencing siRNA or negative control siRNA. Following incubation, all explants were washed with PBS, and luciferase-expressing ATDC5 cells (murine, chondrogenic) in DMEM/F12 1:1 media were cultured on top of the treated cartilage for 24 h at 37 °C. Each well was rinsed with PBS before adding luciferin-containing media (150 ug/mL) and evaluating luminescence by IVIS imaging. The reverse transfection assay was used to compare PEG-DB not functionalized with mAbCII, PEG-DB functionalized with mAbCtrl, commercial reagent lipofectamine 2000, and mAbCII-siNPs. Some treated cartilage plugs for each group were left unwashed to verify gene silencing efficacy of the different formulations independent from the cartilage binding / target capacity. Performance was measured against delivery of the same formulations (targeted and control) loaded with siNEG. Following the measurement of luciferase expression, cell viability of each group was determined using the Promega CellTiter-Glo luminescent cell viability assay following Promega’s standard protocol.

### *In vivo* short-term mechanical loading OA model

C57 mice were mechanically loaded, 3 times per week for 2 weeks. The PTOA model of noninvasive repetitive joint loading was induced by subjecting the knee joints of mice (anesthetized with 3% isoflurane) to 250 cycles of compressive mechanical loading at 9 N. This procedure was repeated three times per week over a period of two weeks using conditions adapted from previous studies (39, 45).

In-joint retention of mAbCII-siNPs in PTOA (2-week loading model) versus healthy mice was also assessed. mAbCII-siNPs were shown to selectively retain in the knee. At 24 h following treatment, the amount remaining in the knee was measured by quantifying fluorescence in the rhodamine-labeled siNPs (**Figure S4**; Exc/Emm: 548/570 nm).

**Figure S4.**
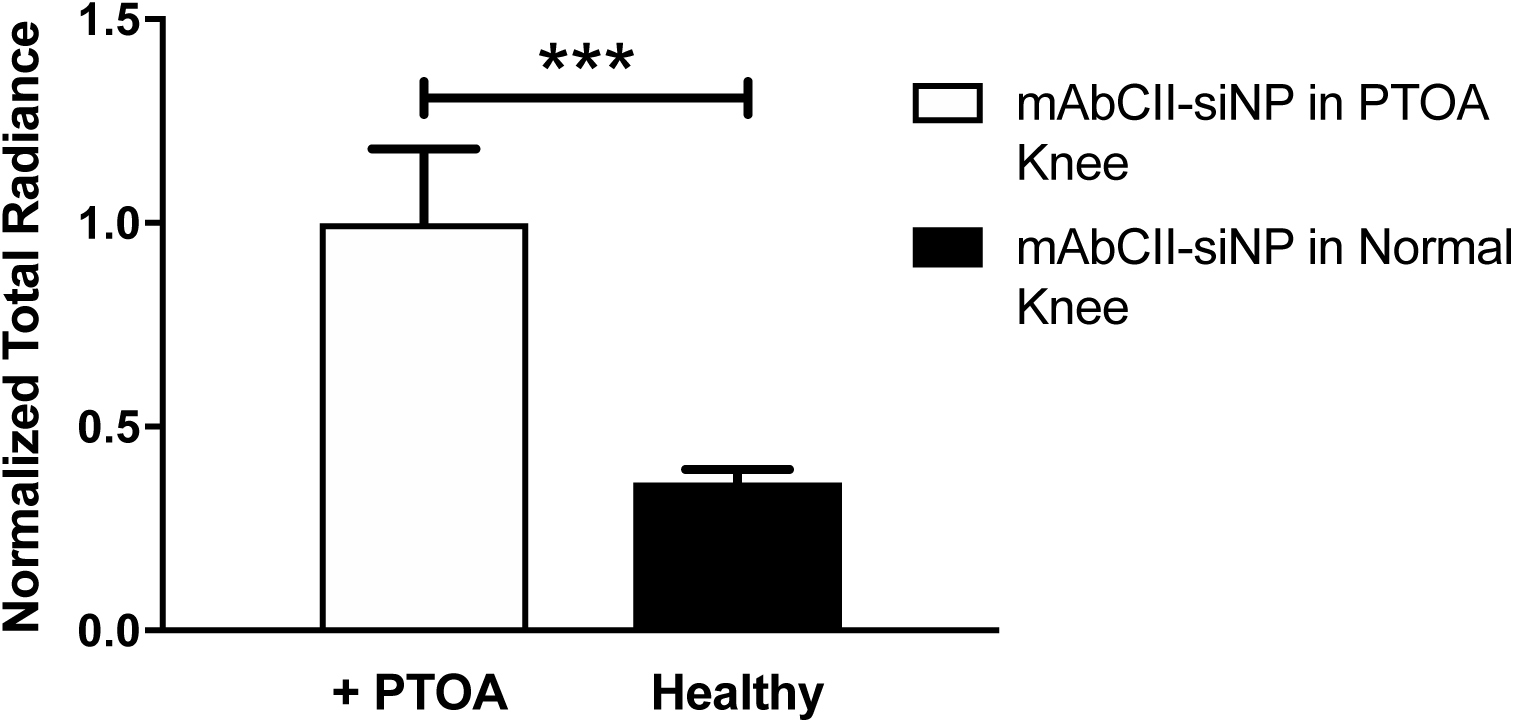
In-joint retention of mAbCII-siNPs in healthy and PTOA mice as measured by rhodamine in siNP polymer (* = p < 0.05; ** = p < 0.01; *** = p < 0.001). Statistics performed with an unpaired t test.

### *In vivo* long-term mechanical loading therapeutic efficacy testing OA model

Mice were aged to 6 months and then subjected to a more rigorous cyclic mechanical loading protocol of 9 N, 500 cycles, 5 times per week, for 6 weeks (48). Doses of siRNA (0.5 mg/kg) were administered into each knee weekly, starting concurrently with mechanical loading. MMPSense and Alexafluor-labeled mAbCII antibody were injected intravenously 24 h before sacrifice to gauge total MMP activity and quantify cartilage damage, respectively.

A high, effective methylprednisolone dose in rodents of 4 mg/kg was administered by intraarticular injection a single time at the beginning of the 6-week time course, which is the duration of the therapeutic study employed with the other groups and the shortest recommended time a patient should wait before repeating intraarticular steroid injection (76, 77). The dose was prepared by initially dissolving in DMSO, diluting in water, and injecting at a volume of 20 μL per knee (60). A separate cohort underwent the same mechanical loading with weekly doses of the same dose of methylprednisolone as a control in direct comparison with the weekly administration of siMMP13 that was speculated to better accommodate the aggressive model, even though less clinically relevant.

### Histology and Immunohistochemistry

Stifles were fixed in 10% neutral buffered formalin and decalcified in Immunocal (StatLab, McKinney, TX). Tissue handling for histopathology was primarily performed in the Vanderbilt Translational Pathology Shared Resource by certified histotechnicians. Fixed tissues were routinely processed using a standard 8 h processing cycle of graded alcohols, xylenes, and paraffin wax, embedded and sectioned at 5 microns, floated on a water bath, and mounted on positively charged glass slides. Hematoxylin and eosin (H&E) staining was performed on the Gemini autostainer (Thermo Fisher Scientific, Waltham, MA). Safranin O staining was performed by hand using a kit (StatLab).

Stifle joints were evaluated by H&E and safranin O in at least 2 serial mid-frontal sections. This histopathologic interpretation was conducted by a board-certified veterinary pathologist under blinded conditions (78). OARSI scores (0-6 semiquantitative scale) were provided for the medial tibial plateau and lateral tibial plateau (54). Simultaneously, a generic score (0-4 semiquantitative scale) was assigned based on safranin O staining of the tibial plateau (**Table S2A**) and on H&E features of degenerative joint disease severity, as defined by cartilage degeneration, meniscal metaplasia, subchondral osteosclerosis, synovial hyperplasia and inflammation, and osteophyte formation (**Table S2B**) (79).

Immunohistochemical staining was performed on a Leica Bond-Max autostainer (Leica Biosystems Inc., Buffalo Grove, IL). All steps besides dehydration, clearing, and coverslipping were performed on the Bond-Max where all the slides were deparaffinized. Heat-induced antigen retrieval was performed on the Bond-Max using Epitope Retrieval 1 solution (Leica Biosystems Inc.) for 20 minutes. Slides were incubated with anti-MMP13 (cat# Ab39012, Abcam, Cambridge, MA) for 1 hour at a 1:750 dilution. The Bond Polymer Refine detection system (Leica Biosystems Inc.) was used for visualization. Slides were then dehydrated, cleared, and cover slipped.

**Table S2.** (**A**) Description of OARSI scale scoring for Safranin O slides of the tibial plateau cartilage (left); (**B**) criteria for scoring of H&E slides to assess overall joint by the degenerative joint disease scale (right).

| Severity Score | OARSI Scale | Degenerative Joint Disease Scale |
| --- | --- | --- |
| 0 | Normal | <u>Within normal limits:</u> mild DJD as a feature of expected age-related change (e.g. attenuation of articular cartilage and proteoglycan loss) |
| 1 | Loss of SO staining, articular cartilage thinning; no defects | <u>Moderate:</u> articular cartilage degeneration and beginning of secondary pathology, synovitis, joint capsule fibrosis, meniscal metaplasia |
| 2 | 1 + fibrillation or pyknotic articular chondrocytes | <u>Marked:</u> metaplasia and/or fragmentation of one meniscus, osteophyte formation, synovitis/hyperplasia |
| 3 | 2 + loss of articular cartilage <50% (e.g. erosion, flap, or callus) | <u>Severe:</u> metaplasia and/or fragmentation of both menisci, total loss of at least 1 articular surface (eburnation), osteophyte formation, advanced synovitis/hyperplasia |
| 4 | 3 + fragmentation and fissuring in an area | <u>Extreme:</u> severe changes plus collapse of stifle joint space and epiphyseal osteolysis |
| 5 | 4 + fragmentation and fissuring in >75% of the cartilage surface | N/A |
| 6 | Total loss of normal articular cartilage (end-stage) | N/A |

### Statistical methods

Data are displayed as mean plus standard error. Statistical tests employed either one-way ANOVA with multiple comparisons test or two-tailed student’s T-test between only two groups with α= 0.05.

### Ethics statement

All animal experiments described herein were carried out according to protocols approved by Vanderbilt University’s Institutional Animal Care and Use Committee, and all studies followed the National Institutes of Health’s guidelines for the care and use of laboratory animals.

